# *M. tuberculosis* microvariation is common and is associated with transmission: analysis of three years prospective universal sequencing in England

**DOI:** 10.1101/681502

**Authors:** David Wyllie, Trien Do, Richard Myers, Vlad Nikolayevskyy, Derrick Crook, Eliza Alexander, Esther Robinson, A Sarah Walker, Colin Campbell, E. Grace Smith

## Abstract

**Background:** The prevalence, association with disease status, and public health impact of infection with mixtures of *M. tuberculosis* strains is unclear, in part due to limitations of existing methods for detecting mixed infections.

**Methods:** We developed an algorithm to identify mixtures of *M. tuberculosis* strains using next generation sequencing data, assessing performance using simulated sequences. We identified mixed *M. tuberculosis* strains when there was at least one mixed nucleotide position, and where both the mixture’s components were present in similar isolates from other individuals. We determined risk factors for mixed infection among isolations of *M. tuberculosis* in England using logistic regression. We used survival analyses to assess the association between mixed infection and putative transmission.

**Findings:** 6,560 isolations of TB were successfully sequenced in England 2016-2018. Of 3,691 (56%) specimens for which similar sequences had been isolated from at least two other individuals, 341 (9.2%) were mixed. Infection with lineages other than Lineage 4 were associated with mixed infection. Among the 1,823 individuals with pulmonary infection with Lineage 4 *M. tuberculosis*, mixed infection was associated with significantly increased risk of subsequent isolation of closely related organisms from a different individual (HR 1.43, 95% CI 1.05,1.94), indicative of transmission.

**Interpretation:** Mixtures of transmissible strains occur in at least 5% of tuberculosis infections in England; when present in pulmonary disease, such mixtures are associated with an increased risk of tuberculosis transmission.

**Funding:** Public Health England; NIHR Health Protection Research Unit Oxford; European Union.

**Research in Context:** 

**Evidence Before This Study:** We searched Pubmed using the search terms ‘tuberculosis’ and ‘mixed’ or ‘mixture’ for English Language articles published up to 1 April 2019. Studies, most performed without the benefit of genomic sequencing, report mixed TB infection from a range of medium and high prevalence areas and show it to be associated with delayed treatment response. Modelling suggests detection and treatment of mixed TB infection is an important goal for TB eradication campaigns. Although routine DNA sequencing of *M. tuberculosis* isolates is becoming widespread, efficient methods for detecting mixed infection from such data are underdeveloped, and the true prevalence of mixed infection and its association with transmission is unclear.

**Added Value of This Study:** This study investigated a large series of TB isolations obtained as part of a routine Mycobacterial sequencing program by two reference laboratories, in a low incidence area, England. We developed an efficient generalisable approach to identify transmitted mixed *M. tuberculosis* infection; our approach is capable of sensitive and specific detection of a single mixed nucleotide position. We identified mixed infection of similar strains (‘microvariation’) in about 9.2% of the *M. tuberculosis* samples which we were able to assess, and found evidence of increased transmission from individuals with mixed infection.

**Implications of All the Available Evidence:** TB microvariation is a risk factor for TB transmission, even in the low incidence area studied. Although an efficient and highly specific technique identifying microvariation exists, it relies on comparison with similar sequences isolated from other patients. Sharing of sequence data from the many TB sequencing programs being deployed globally will increase the sensitivity of microvariation detection, and may assist targeted public health interventions.

## Introduction

The WHO goal of eliminating TB transmission ^1-3^ will be aided by understanding which cases of *M. tuberculosis* infection are most likely to transmit infection, and of transmission networks.^4^ The latter can be discerned from patterns of single nucleotide variation (SNV) derived from next generation sequencing (NGS) of *Mycobacterium tuberculosis* isolates^5^; such networks are being studied as part of an ongoing national TB control program in England. Sequencing data is generated after clinical specimens (e.g. sputum) are cultured in broths, and DNA from positive broths is extracted. The resulting short read sequencing data is mapped to a reference genome, an appropriate strategy given the relatively limited diversity of *Mycobacterium tuberculosis.*^5^

Infections involving mixtures of *Mycobacterium tuberculosis* strains are well described^6-12^. Such infections are currently of clinical interest since response to treatment is known to be delayed in individuals with mixed infection.^10, 13, 14^ Mixed infection may reflect either the development of intra-host diversity during chronic infection^6^, or co-infection by more than one strain circulating in the community in which the affected individual resides.^15, 16^ Co-circulating strains are commonly closely related,^15^ and co-circulation is ongoing in hyperendemic settings, in which mixed isolates are reported in 10-30% of cases.^8-10^ In either the chronic infection or superinfection scenario, mixed infection might be associated with raised transmission risk, although data to support this is not available at present.

Prior to widespread use of NGS, mixed infections were detected by analysis of multiple picks of bacteria growing on solid media, and by analysis of their MIRU-VNTR typing profiles.^7^ Starting from NGS data, there is no consensus on how to detect, quantify and report mixtures. Published NGS-based mixture detection algorithms identify mixed bases at positions reflecting divergence of ancestral TB lineages, which is an effective way of detecting mixtures of distantly related isolates.^17, 18^ A separate approach, potentially applicable to similar samples, involves the enumeration of the number of sites to which multiple different bases are mapped, together with analysis of the frequency of the variants at these sites.^19^

Here, using data from the English prospective TB sequencing programme, we describe practical limitations restricting use of existing mixture detection algorithms, and propose a new approach to identify mixtures of TB strains using NGS data^19^. We show mixed infection occurs in at least 5% of TB isolates in England and is associated with an increased risk of disease transmission.

## Methods

### Isolation of DNA from Mycobacteria and sequencing

This study includes all *M. tuberculosis* isolates sequenced between 01/01/2016 and 15/12/2018 by the two English Mycobacteriology reference laboratories; these receive samples from across the English NHS. One laboratory, covering the Midlands and North of England, performed universal sequencing, initially as part of method validation, from 1 January 2016. The second laboratory covers the south of the country and performed universal sequencing from 7 January 2018. The two sites are managed as a single service, use identical DNA extraction protocols and bioinformatics pathways, but use different Illumina sequencing platforms (MiSeq and HiSeq, respectively; see also Web Appendix).

### Assessment of mixed sequences using base call frequencies

High quality base counts mapped to the reference genome were examined. We excluded positions susceptible to mismapping due to Mycobacterial-non Mycobacterial homology^20, 21^. Mapping software provides estimates of the probability that a read is mapped to the wrong location (mapping quality), something which may create apparent mixtures. Using only reads with high quality mapping (mapper reported estimated probability of mismapping ≤ 10^−3^), we identified positions where the number of minor variant bases observed significantly exceeded the number expected, using a Binomial Test with Bonferroni control for multiple comparisons. We defined such positions as mixed sites (M-sites) if no other apparently mixed sites existed with 10 nt (see Web Appendix for details). Positions which did not meet criteria to confidently call either a Mixed (M) site, or an unmixed, single base (A,C,G,T) were called as uncertain (N-sites). We identified the specimen’s ancestral lineage using the published phylogeny^22^ and assessed inter-lineage mixtures of two, or more than two, samples using F2 and F47 statistics, as described^17^.

### Computation of single nucleotide variant distances between samples (SNV)

We compared sequences pairwise, considering a SNV present if a base (A,C,G,T) was confidently called in one sequence, and a different base confidently called in the other sequence. A SNV was counted at a given position only if the two sequences differed in confident (A,C,G,T) consensus calls; it was not considered to be present if one or both consensus calls were uncertain (N-sites) or mixed (M-sites) (Fig. S1).

### Detection of mixed sequences and the MixPORE algorithm, and software implementations

In this work, we developed a novel algorithm (MixPORE), to efficiently identify mixed sequences from multiple sequencing platforms. This tests the hypothesis that in a given test sequence *s*_*t*_, M-sites occur at significantly higher frequency in those positions P where recent evolution has occurred (positions of recent evolution, PORE), relative to all other positions (P’). Critically, this intra-sample comparison (Fig. S2) is expected to be robust to different intrinsic rates of M-sites, e.g. due to sequencer characteristics. For computational efficiency, PORE were identified operationally without reconstructing phylogenies by comparing *s*_*t*_ with highly similar sequences from different individuals, based on SNV distances, using a pre-specified SNV threshold; M-sites occurring at PORE were termed mixed and recently evolved sites (MRE-sites). The Web Appendix includes more details, including of *findNeighbour3*, an open source bacterial relatedness server which implements MixPORE.

### Risk factors for sequences having mixed bases

To examine risk factors for mixed infection, we applied MixPORE separately to the whole dataset using 100 SNV thresholds (or 5, 20, or 50 SNV thresholds in sensitivity analyses) to detect neighbours for PORE determination. We classified samples as mixed (defined as at least one MRE-site) or not mixed. We classified individuals as having mixed infection if they had at least one mixed sample, and used logistic regression models to relate individual’s mixture status to clinical features ascertained during routine surveillance of TB in England and stored in the PHE Enhanced TB Surveillance System.

### Mixed bases and isolation of subsequent similar samples

To determine whether the detection of mixed bases was associated with TB transmission, we applied MixPORE to each sample of interest *s*_*t*_ in the order samples were received, only identifying neighbours among samples received before *s*_*t*_ and from a different individual. Then, we computed the time to subsequent isolation of a sample closely related to *s*_*t*_ from a different individual, compatible with transmission. We considered samples *closely related* if they were (i) 0-3 SNV of *s*_*t*_ and (ii) were isolated after the first case, using Kaplan-Meier estimates of time to similar isolation, and Cox Proportional Hazards models, censoring individuals with no subsequent isolation at one year. A range of sensitivity analyses were performed around the definition of *closely related*, see Web Appendix.

### Funding Source

This study was supported by the National Institute for Health Research (NIHR) Health Protection Research Unit (NIHR HPRU) in Healthcare Associated Infections and Antimicrobial Resistance at University of Oxford in partnership with Public Health England [HPRU-2012-10041], and by the NIHR Biomedical Research Centre, Oxford. VN received funding from the European Centre for Disease Control under Grants 2014/001 and 2018/001. TEAP, DCW and ASW are NIHR Senior Investigators. The views expressed are those of the author(s) and not necessarily those of the NHS, the NIHR, the Department of Health or Public Health England. The sponsors of the study had no role in study design, data collection, data analysis, data interpretation, or writing of the report. The corresponding author had full access to all the data in the study and had final responsibility for the decision to submit for publication.

### Ethical framework

Tuberculosis surveillance is one of the Public Health England’s key roles. The Health and Social Care Act 2012 and related statutory instruments provide a legal basis for the activity. This work is part of service development carried out under this framework, and as such explicit ethical approval is unnecessary.

## Results

### Mixed *M. tuberculosis* infection in severe infection: a motivating example

In England, groups of *M. tuberculosis* isolates which differ by low SNV distances are now routinely reviewed to direct public health interventions. For example, we noted the isolation of six similar *M. tuberculosis* strains from samples taken within a month of each other. Isolates A-C are from patient-1, who subsequently died of disseminated tuberculosis, D and E from respiratory samples from patient-2, while isolate-F is from a respiratory sample from patient-3 (Table 1). High confidence base calls across the 4.4 million base genome differ between these six samples at only sixteen positions, with A and B (from patient-1) differing at 12 positions; isolates identical to A and B were obtained from patient-2 and patient-3.

**Table 1.**
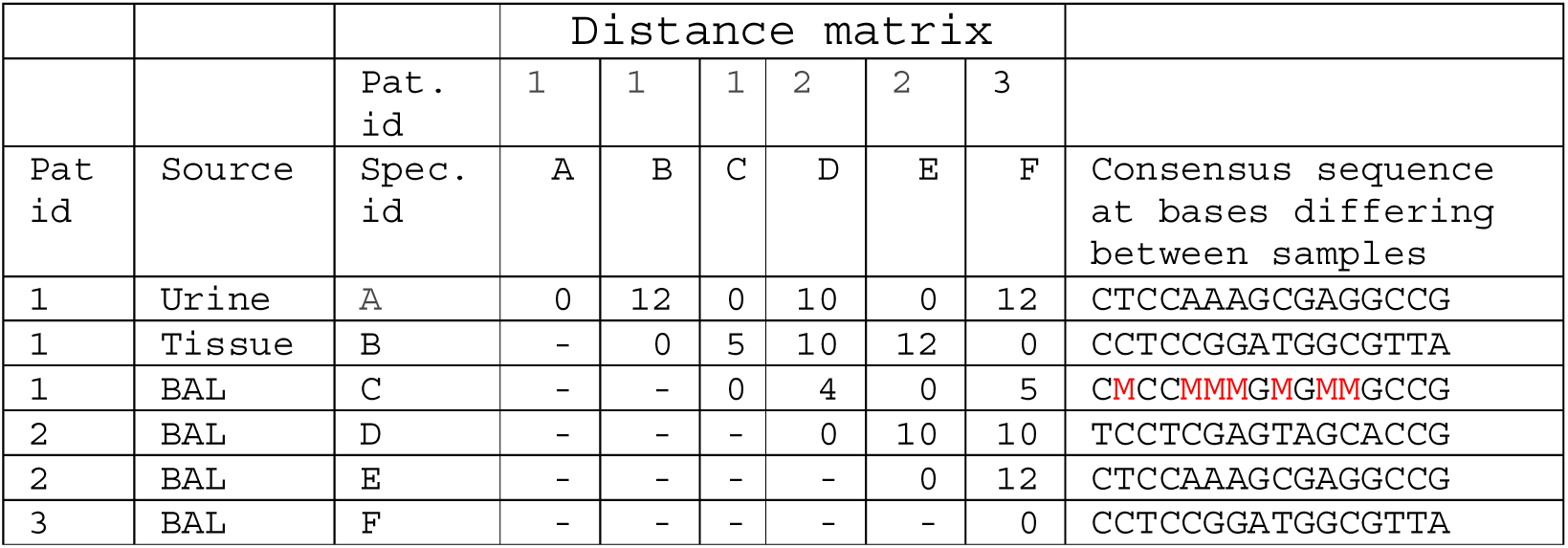
Single nucleotide variation, and sequences, of six samples. Six samples from three patients, all taken within a month of each other, differed in consensus sequence at sixteen of the 4.4 million positions analysed. Bases which are mixed (M-sites), assessed using a binomial test comparing observed with expected minor variant frequencies in mapped sequence, are shown as M. For bases which are not mixed, the consensus base call is shown. Bases failing quality cutoffs were marked N, but none exist at the positions shown.

Examining the bases mapped to each site using per-base binomial testing (see Methods) shows sample-C from patient-1 contains multiple mixed positions (M-sites) at the sites differing between A and B. This kind of within-patient heterogeneity, which is reported to be associated with impaired treatment response,^10, 13, 14^ is not evident using standard methods of consensus base calling, using which C appears identical to (that is, has a zero pairwise SNV distance from) sample A. Consequently, we decided to develop automated methods detecting mixed TB infection using our routinely sequenced data.

### Estimates of mixed base numbers differ markedly depending on sequencing technology

Between 1 January 2016 and 15 December 2018, 7,251 *M. tuberculosis* samples were sequenced in the two Reference laboratories in England (Figure 1). Of these, 6,560 samples (90%) passed quality control; 401 samples were excluded due to insufficient read depth, and a further 290 were excluded as they appeared to consist of mixtures of more than two strains, as evidenced by an elevated F47 quality statistic, which we have observed is associated with breakdowns in laboratory processing.^17^ 4,195 samples were sequenced using Miseq, and 2,385 using Hiseq; they derived from 4,928 individuals. The number of bases detected as mixed (M-sites) differed markedly between the two centres, with a median of 7 *vs.* 99 bases, respectively in the centre using HiSeq *vs.* MiSeq technology (*p* < 10^−8^, Fig. S3), whereas differences in the number of otherwise uncertain (N-sites) were much smaller (Fig. S3).

**Figure 1.**
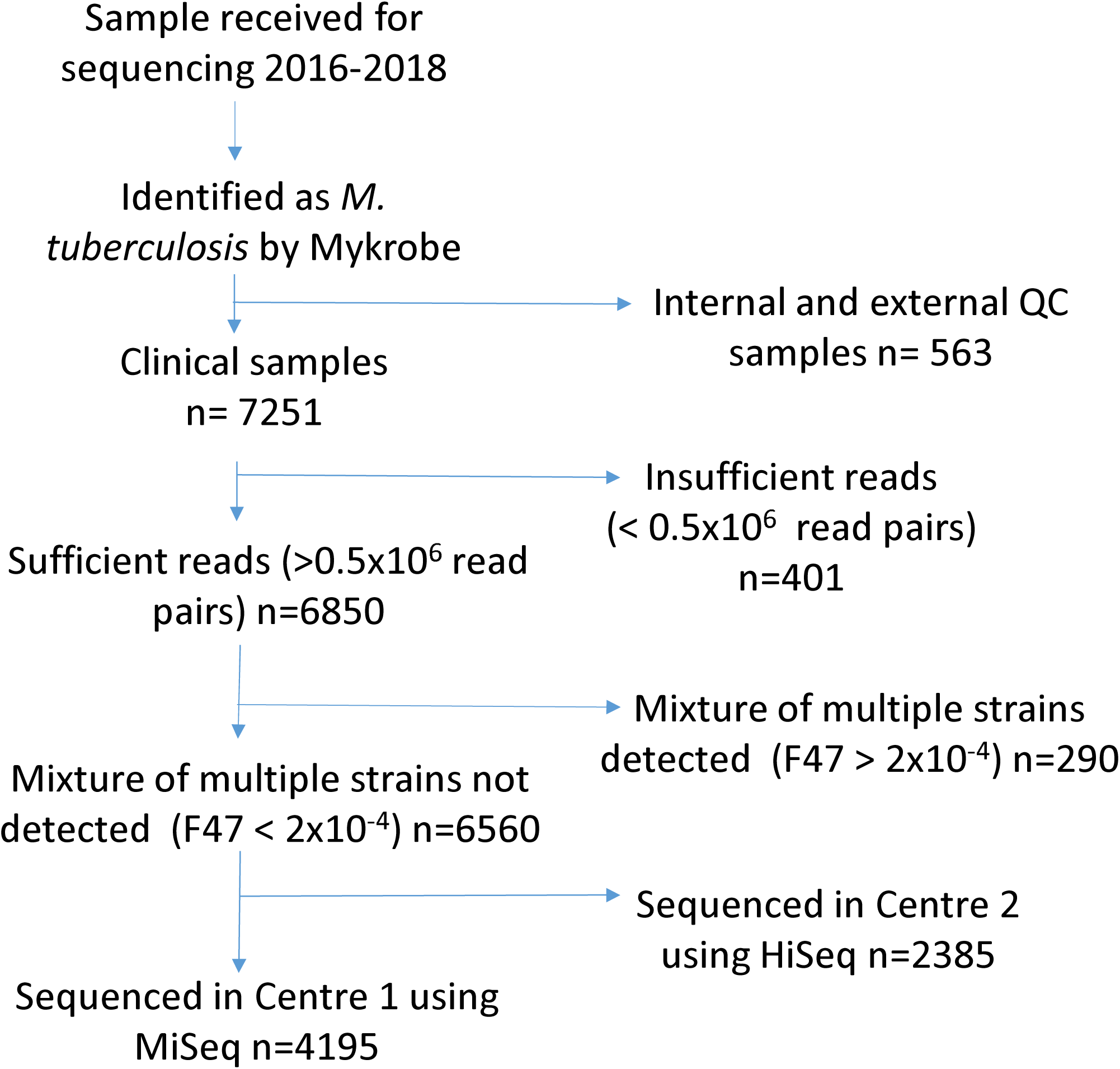
Clinical samples examined. A flow chart showing clinical sample selection.

To understand how sequencing technology affects the number of M-sites detected, we sequenced ten Mycobacterial DNA preparations, which formed part of a quality control distribution, using three different Illumina sequencing instruments. All samples had a population of M-sites detected (Figure 2), a proportion of which occurred in clusters being grouped together (Figure 2, yellow bars). The number of M-sites, the proportion of M-sites which are clustered, and the mean minor variant frequency within an M-site, all differed markedly between sequencing technologies (Kruskal-Wallis, p < 10^−4^ for all three comparisons; Figure 2, Figure S4). One sample, 7, was a mixture of two Lineage-4 strains, as judged by analysis of variation at positions determining ancestral phylogeny: the F2 statistic,^17^ which reflects the proportion of the less common lineage in the mixture, was between 0.40 and 0.44 for all three sequencing technologies. In consequence, we would expect to, and did, observe several hundred bases to be mixed in this sample,^22^ representing the positions of difference relative to the common ancestor of the two strains (Figure 2, arrow).

**Figure 2.**
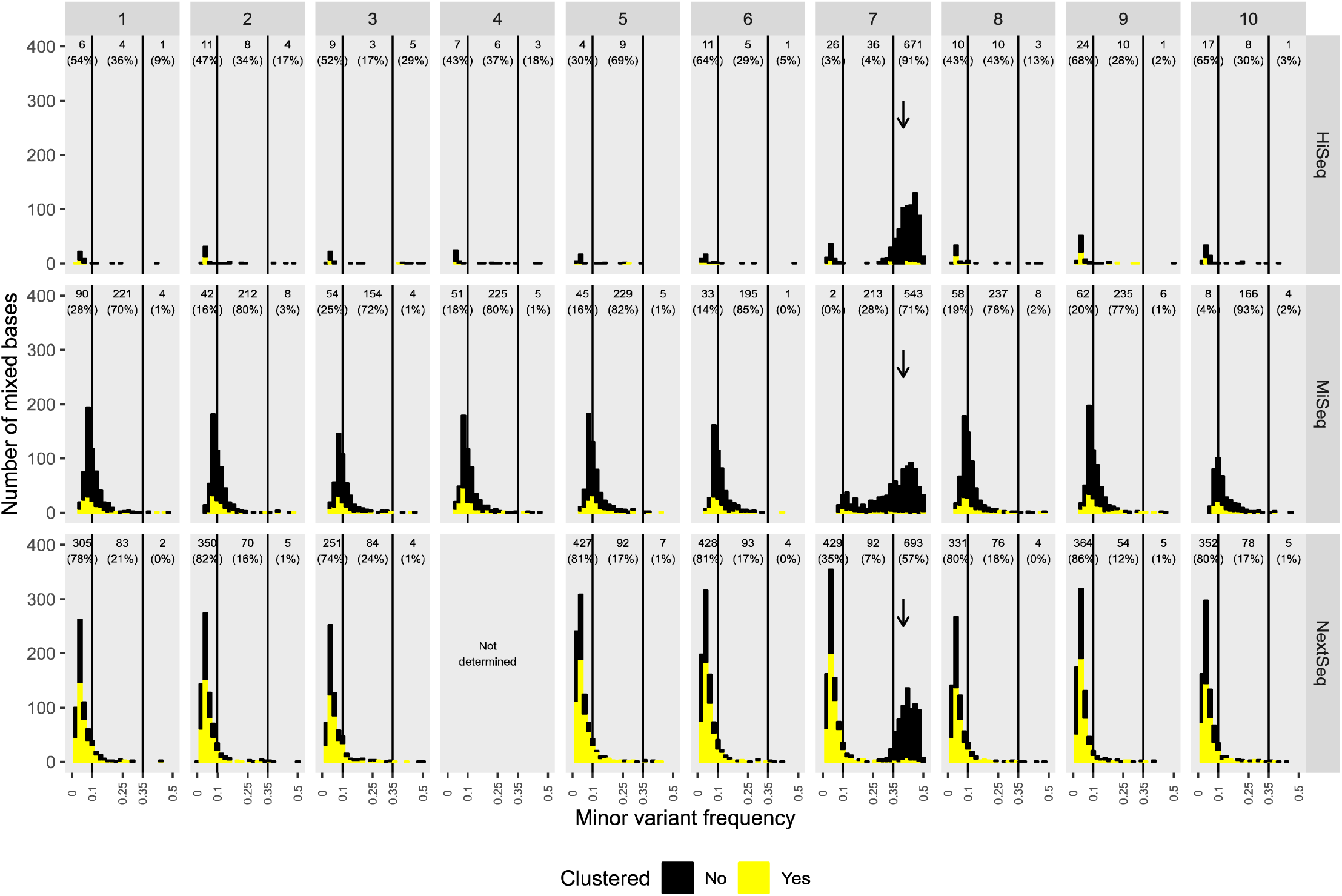
Detection of mixed bases is influenced by sequencing technology. Histograms depict numbers of mixed bases, and the minor variant frequency, for ten samples sequenced using three different technologies. Yellow bars reflect the number of “clustered” mixed positions, defined as mixed positions within 10 nucleotides of another. Vertical lines are at minor variant frequencies of 0.1 and 0.35, which are arbitrary positions. Arrows reflect a population of mixed bases associated with a mixed sample (sample 7).

### Statistical testing for mixed sequences

For public health purposes, mixture detection algorithms that critically depend on a single sequencing technology are undesirable, as the ability to analyse data from multiple laboratories and jurisdictions is essential.^5, 23^ We therefore developed an algorithm, MixPORE, which compares the number of M-sites located within positions of recent evolution, designated MRE-sites, with the number of M-sites located in other parts of the genome (see Methods, and Fig S2D). The rationale is that positions of recent evolution (PORE) are likely to contain mixtures of biological, rather than technical, origin given what is known about co-transmission of similar TB isolates,^15^ while the intra-sample comparison of PORE *vs*. other sites controls for technical causes of variations in M-site calling.

First, we assessed the performance of MixPORE using simulated phylogenies, in which fifty samples evolved from a single ancestor, and in which one mixed sample exists (see Web Appendix). These simulations indicate that MixPORE has near perfect sensitivity and specificity when determining mixtures in samples with two or more close neighbours (Fig. S5); for other samples, MixPORE cannot be applied. As expected, simulations indicate sensitivity of mixture detection by MixPORE is not strongly influenced by MiSeq *vs.* HiSeq sequencing platform (Fig. S6).

### Performance of MixPORE Statistical testing for mixed sequences

We applied the MixPORE algorithm to the 6,560 consecutive *M. tuberculosis* samples passing quality control (Figure 1). In our main analysis, we assessed PORE by comparing sequences with those differing by less than 100 SNVs; this corresponds to the samples having a common ancestor less than about 150 years ago, or in the mid-Victorian era.^5^ 3,690 samples (56.3% and 55.6% of samples sequenced by HiSeq and MiSeq, respectively) had similar samples from least two different individuals, and so were assessable by MixPORE (Figure 3). Of these 3,690 assessable samples, 341 (9.2%) of samples had at least one MRE-site detected, so at least 341/6,560 (5.2%) of English TB isolates have this kind of variation. Only 9 had more than 10 MRE-sites, and these nine were all mixtures of TB isolates from different deep branches of the ancestral TB phylogeny, as judged by elevated F2 statistics;^17^ of these, all but one had over 50 MRE-sites detected (Figure 3).

**Figure 3.**
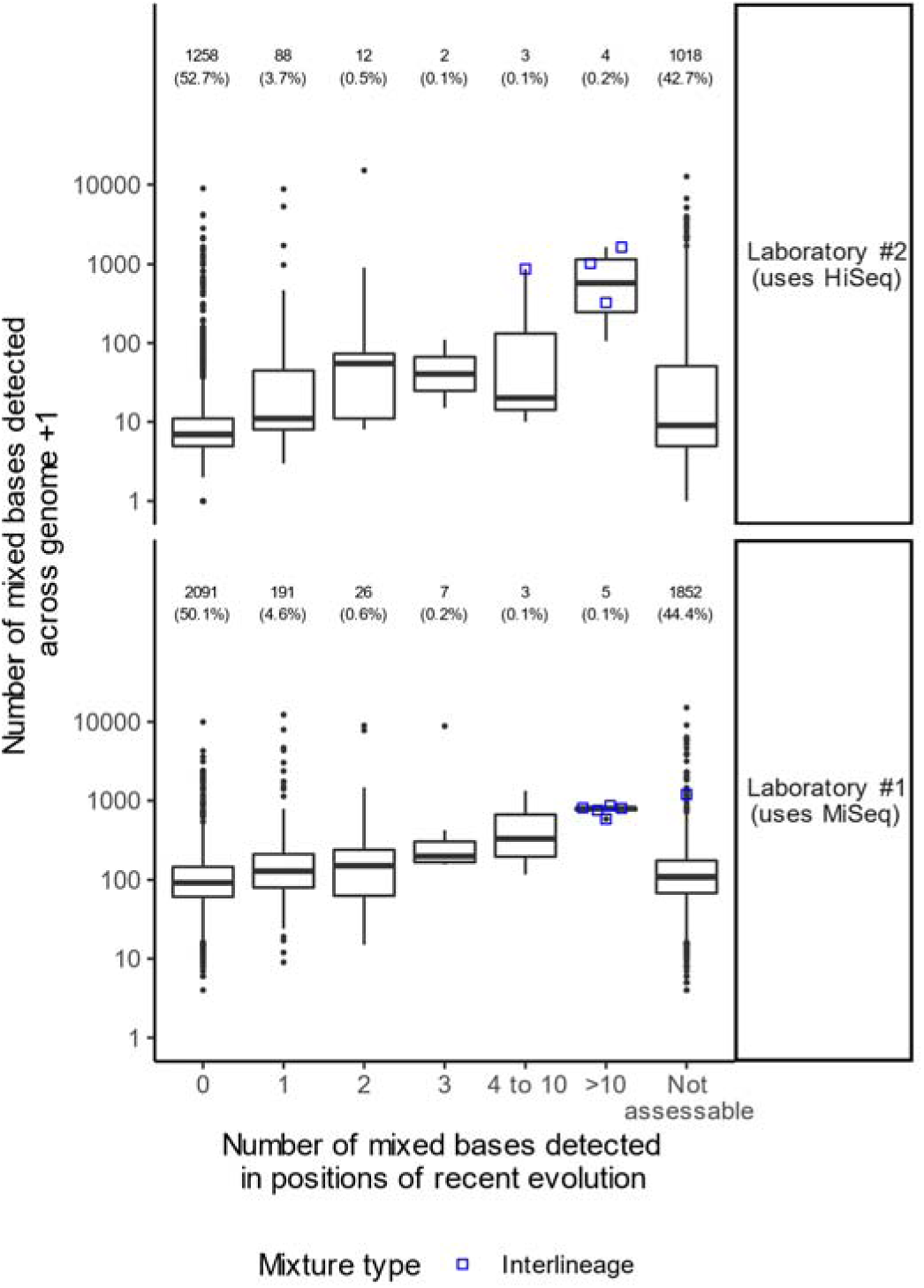
Relationship between total mixed base detection and that by MixPOR. Relationship between the numbers of mixed bases detected using per-base examination (y-axis) and by MixPORE (x-axis). The analysis is stratified according to whether the sample was analysed by MiSeq or HiSeq technology. Numbers show number (percent of total for MiSeq/HiSeq) of samples in each group. Large circles indicate inter-lineage mixtures, detected by F2 statistic computation as described^17^. This figure shows results when a SNV of 100 is used to select neighbours for the MixPORE algorithm.

We considered whether, as in simulated sequences (Fig. S5), MixPORE detects mixed bases in a highly sensitive manner in real data. Were this the case, for every mixed base detected by MixPORE (MRE-site), we would expect an increase of exactly one in the number of mixed bases detected by per-base binomial testing (M-site). We tested this by modelling the relationship between these two quantities (Fig. 3), using quantile regression. For each increase in MRE-sites, M-sites increased by median (±standard error of median) 1.13 (±0.12) and 1.17 (±0.12) for HiSeq and MiSeq samples, respectively. A similar relationship was seen when other thresholds were used to identify PORE (Figs. S7, 8, 9).

### Properties of individuals without close neighbours, who cannot be assessed

Since MixPORE can only operate if similar samples from three different individuals can be identified, we investigated the characteristics of the individuals on whose samples the algorithm could *vs*. could not be applied (Fig. 4A). The individuals without any assessable samples (2,240/4,928 (45%)) were independently less likely, compared to those who were assessable, to be UK born (aOR=0.55 vs born elsewhere, 95% CI 0.47,0.64), to have a known history of imprisonment (aOR=0.59, 95% CI 0.42,0.83), and to have pulmonary disease (aOR=0.82 vs no pulmonary disease, 95% CI 0.72, 0.94), the latter two being known to enhance transmission. Unasssessable individuals were also more likely to be older (e.g. aOR=2. 19, 95% CI 1.61, 2.92 for over 55 relative to 18 years or younger), and to be infected with TB lineages 1 or 3 (e.g. lineage-1 *vs.* lineage-4, which is commonly isolated in the UK, aOR=5.89, 95% CI 4.66,7.26). Sensitivity analyses using a range SNV thresholds all yielded similar conclusions (Supp. Data D1).

**Figure 4.**
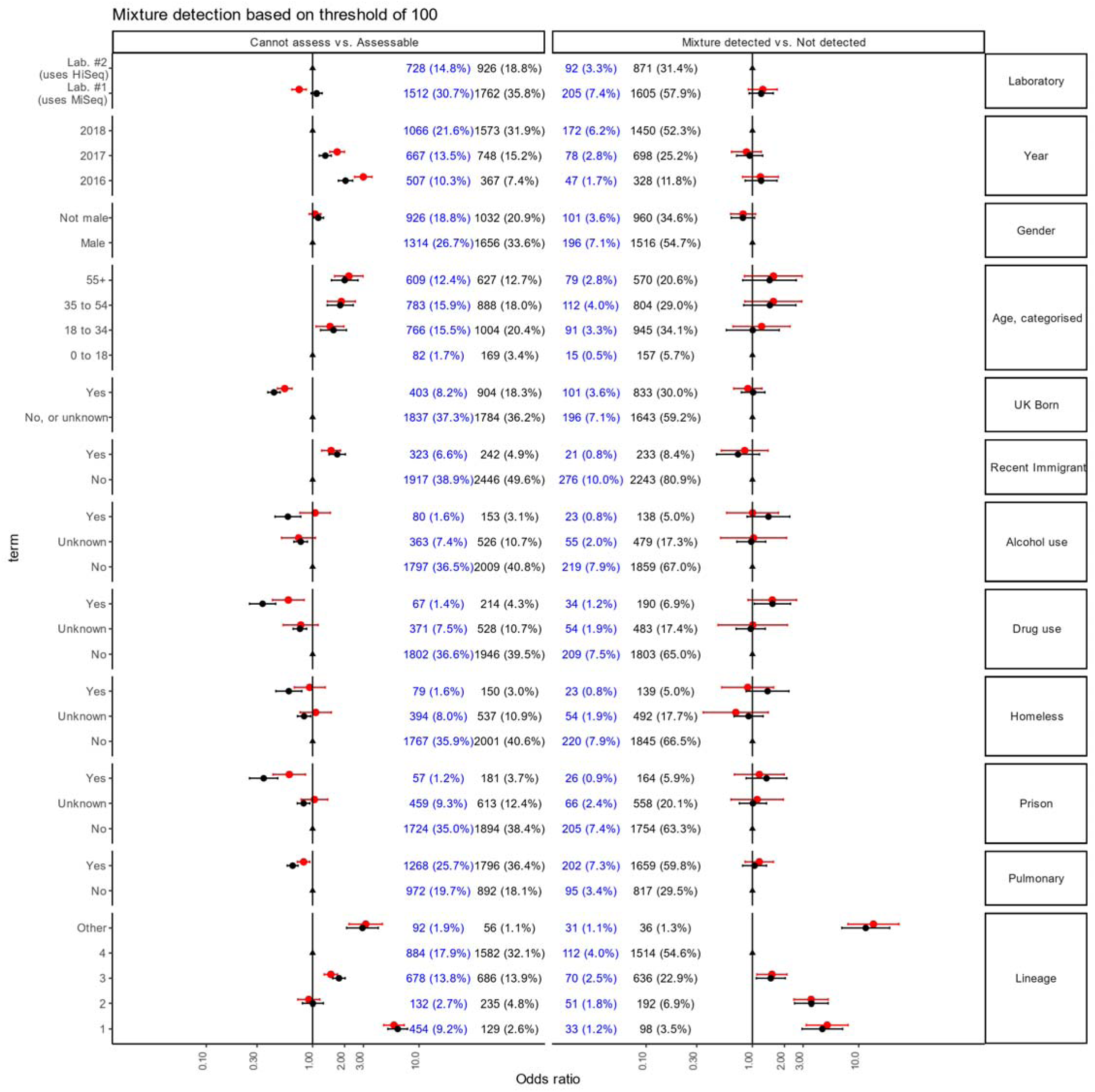
Characteristics of individuals with and without mixed infection. The MixPORE algorithm, which requires two or more samples from different individuals to be present within a 100 SNV threshold, was applied to all samples. Mixed samples were defined as the presence of at least one mixed base. We classified individuals as having mixed infection if they had at least one mixed sample. Panel A shows odds ratios having a sample assessable by MixPORE among all 5,013 individuals studied. Blue numbers (% total patients) are number of individuals assessable *vs* not assessable (black numbers). Panel B shows odds ratios for mixed infection among the 2,865 individuals from whom an assessable sample was derived; blue numbers (% total patients assessable) refer to patients with mixed infection vs. those without. Black dots and error bars are odds ratios and 95% CI from univariable logistic regression; red symbols are derived from multivariable logistic regression including all variables shown.

### Risk factors for mixed infection

Among the individuals with samples which could be assessed, isolation of bacteria which were not of lineage-4 (1,080 (40%) vs 1,626 individuals; *vs.* lineage-4, for lineage-1 aOR=5.05 95% CI 3.21,7.93; for lineage-2 aOR 3.57, 95% CI 2.46,5.16) was the only significant risk factor.

### Mixed infection is associated with the subsequent isolation of similar isolates

Finally, we considered whether mixed infection was associated with transmission. For isolates from patients with pulmonary disease (selected as they are potentially infectious) we computed the time to the first subsequent isolation of closely related (≤3 SNV, see Methods) isolate(s), from any body site, from a different patient, used as a measure of transmission. We examined each isolate in the order it was obtained, assessing mixture status using MixPORE applied to preceding samples only. Since different bacterial lineages display different rates of mixed samples (Fig. 4), as well as different biological behaviours,^24^ we conducted separate analyses for the most common lineage circulating in the UK, lineage-4 and for other lineages (Fig. 5A, B). In lineage-4 pulmonary isolates, mixed samples were associated with increased likelihood of subsequent isolation (*p*=0.02 by log-rank Test; hazard ratio=1.43, 95% CI 1.05,1.94, from Cox Proportional Hazards model). Such an effect was not evident in other lineages (Fig. 5B) in this dataset. To control for possible over-ascertainment of similar cases by mixed samples, we analysed the subgroup of samples with zero or one mixed base, measuring time to subsequent isolation of samples zero or one SNV different for samples without mixed bases, and zero SNV different for those with one mixed based. This analysis gave essentially identical results to those in Fig. 5A (*p*=0.02 by log-rank test).

**Figure 5.**
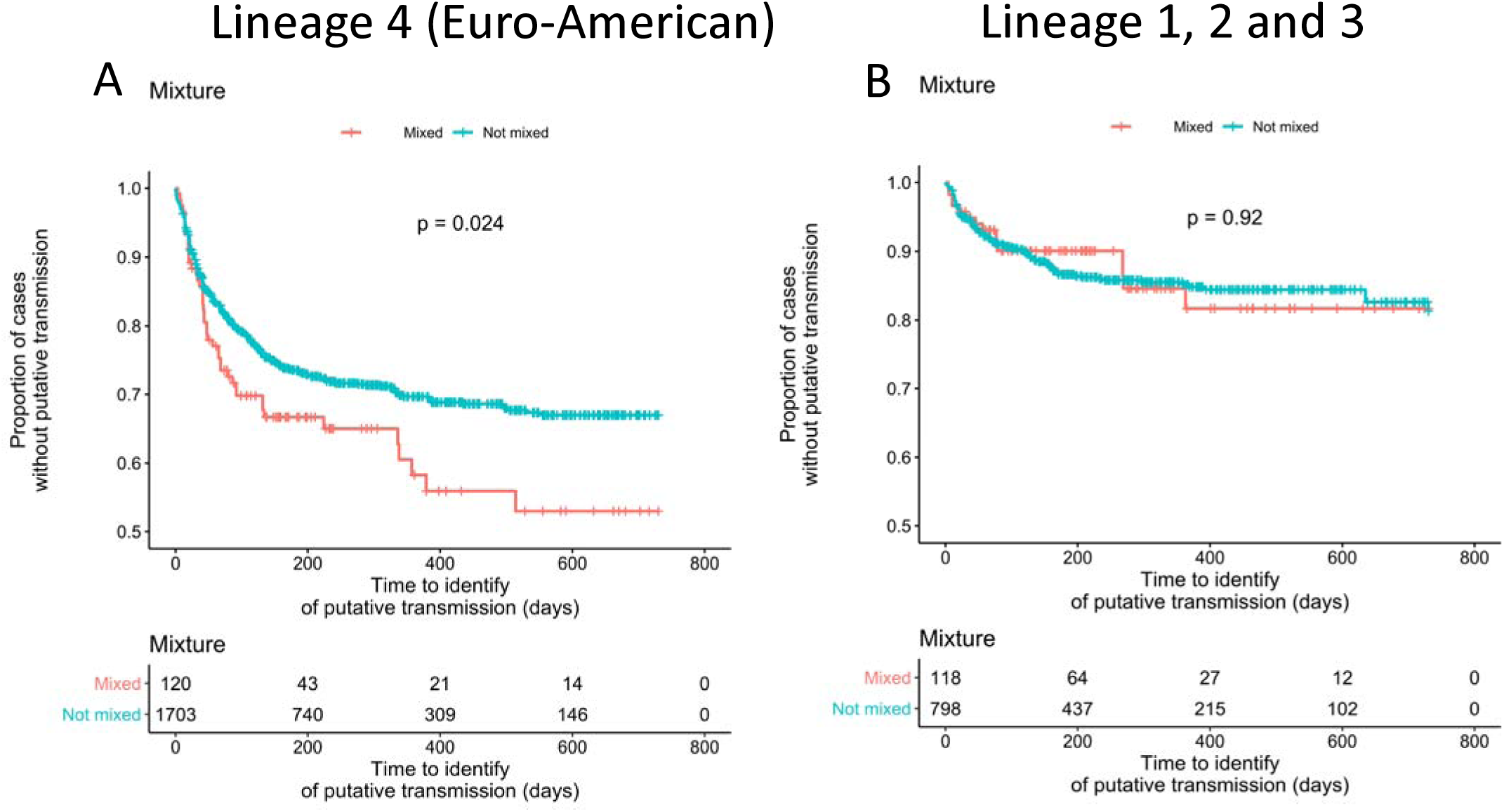
Mixed base detection and subsequent isolation of similar samples. Kaplan Meier survival curves estimating the time between isolation of a sample of interest, and the subsequent isolation of a similar isolate from a different individual. Analyses are stratified by lineage-4 vs other lineages, by whether at least one mixed base is detected by MixPORE. *p-*values are from Log-Rank tests.

## Discussion

Mixed TB infection commonly involves the co-infection of an individual with similar isolates co-circulating in a community,^15^ and/or the generation and subsequent transmission of diverse strains from a chronically infected host. In these situations, the clones making up the mixed infection differ at a small number of positions only, representing divergence from the common ancestor of the mixture.^5^ Here, we used a novel and specific algorithm which, both simulations and studies on real data indicate, is likely to capture a large proportion of the existing transmitted variation in cases where it can be applied. Among the ∼ 56% of samples which could be assessed, transmitted microvariation affected a small number of bases: only about 0.7% of samples (∼ 1 sample in 150) had four or more mixed bases detected. By contrast, 9.2% of samples had at least one mixed base. Thus, we can place a lower limit of about 5% (∼ 1 in 20) of UK isolates having transmitted microvariation. We determined that the microvariation we detected, even if present in only a single position, was biologically relevant: for cases with pulmonary lineage-4 disease, microvariation was associated with significantly shorter time to subsequent isolation of closely related isolates (Fig. 5A).

Strategies for detecting mixed TB infection^6-12^ are not well established, and the approach described is a new approach in this area. Other approaches to addressing the same problem have used retrospective sample collections analysed on an Illumina HiSeq platform; per-base minor variant frequencies, and the number of apparently mixed bases, were analysed using Bayesian techniques^19^. The small number of mixed bases detected in our approach (typically one or two) was below a threshold used in this work; additionally, its application to our data is problematic since both the number of bases with high minor variant frequencies, and the minor variant frequency distribution, is strongly influenced by the type of Illumina sequencer used (Figure 2), an important but previously unappreciated complication of drawing public health inference from sequencing using differing technologies.

Our study has a number of limitations. Firstly, we used culture to amplify *M. tuberculosis* isolates, which may reduce diversity,^25^ and so may underestimate the importance of microvariation in TB infection. The introduction of culture independent sequencing methods^26^ may address this. Secondly, although we observed microvariation to be more commonly detected in lineages 1-3 than in the most common UK lineage-4, the biological basis for this needs further study. Thirdly, although mixed infection has been associated with delayed response to therapy in high incidence settings,^10^ we were unable to determine whether this occurs in the UK as the UK’s national TB surveillance scheme does not collect suitable data on treatment response; it records only rates of treatment completion within 12 months of diagnosis, or within 24 months for cases with rifampicin or multidrug resistance.

Finally, the technique detects potentially transmitted microvariation, and requires sequenced isolates from at least three different individuals to determine this. In our work, about 45% of cases were not assessable; as might be expected, they were less likely to have recognised risk factors for UK-based transmission,^27^ more likely to have non-pulmonary disease, to be older (perhaps reflecting reactivation of latent disease), and more likely to be infected with lineages which are predominantly imported into the UK.^27^ The proportion of unassessable cases is expected to fall, as we have only one year’s data from London and the South East where about two thirds of TB cases in England occur. At present, about 70% of cases of pulmonary disease are culture positive,^27^ and all are being sequences, so relatively complete coverage of the TB populations transmitted in England is accruing.

Overall, this work establishes *M. tuberculosis* sequence microvariation as a risk factor for TB transmission. Increasing numbers of countries are routinely sequencing *M. tuberculosis*; this, together the approach described here, will allow a better understanding of the region(s) of the genome where microvariation contributes to disease transmission, disease progression and drug response, as well as the identification of individuals at high risk of transmitting TB.

## Supporting information

Web appendix

## Data availability

The data analysed is available at https://github.com/davidhwyllie/findNeighbour3/blob/master/doc/demos.md.

## Author Contributions

Study design: DW, CC, TD; Software development: DW, TD; Performed experiments: EA,GS,ER,VN; Performed analyses: DW; Wrote first draft: DW, TP; Revised manuscript: TP, ASW. Critical review of manuscript: all authors.

