## Supplementary material for "*M. tuberculosis* microvariation is common and is associated with transmission: analysis of three years prospective universal sequencing in England": Web appendix

### Supplementary Methods

#### Isolation of DNA from Mycobacteria and sequencing

This study includes all Mycobacteria processed between 01/01/2016 and 15/12/2018 by Public Health England Mycobacteriology reference laboratories. There are two such laboratories; one, covering the Midlands and North of England (~ 15 million person catchment) uses MiSeq sequencing technology as described^1^, and, after a run-in period, was fully operational by 1 January 2016. The second covers the south of the country (~ 35 million person catchment) and uses HiSeq sequencing. It became fully operational on 7 January 2018. When sequencing was occurring, all positive Mycobacterial isolates received in the reference laboratory were processed through the same pathway. Briefly, clinical specimens were decontaminated and inoculated into Mycobacterial Growth Indicator Tubes (MGIT) tubes, and DNA extract from positive broth cultures was sequenced as described in Methods, and ref ^1^.

#### Routine bioinformatic processing

The routine bioinformatics pipeline deployed by Public Health England has been previously described^2^ (see also Software Implementation section). Briefly, specimens identified as containing *M. tuberculosis* based on a k-mer analysis using Mykrobe^3^ were further processed,^1^ reads comprising high quality read (Illumina read quality phred score q >=30) were mapped to the H37Rv v2 genome (Genbank:NC_000962.2)^4^, and VCF files generated using Samtools mPileup, with additional basecalling using GATK VariantAnnotator v2.1.

#### Assessment of mixed sequences using base call frequencies

High quality base counts, as identified by SamTools, were extracted from mapped data. We defined the major variant as the most common base at each position. We expect variants other than the major variant to be present at a frequency of 0.001 or less, a figure corresponding to the Q30 mapping error estimate provided by the mapper, which was used to select high quality mapping reads. Candidate positions with mixtures, defined as positions with elevated non-major variant counts relative to this expectation, were identified using a binomial test using a *p v*alue of 2.3x10^-9^, corresponding to *p* < 0.01 with Bonferroni correction for multiple comparisons across the genome. We observed close clustering of candidate positions with mixtures in some genomic regions, something we considered likely to reflect technical factors (see Results Fig. 1), and considered these bases mixed (‘M’ sites) only if no other apparently mixed sites existed with 10 nt. Candidate positions with mixtures within 10nt of another candidate were considered uncertain (‘N’ sites).

#### Identification of bases for which we cannot be confident of the sequence

We did not analyse the genomic regions which are susceptible to mismapping due to Mycobacterial-non Mycobacterial homology.^5^ These regions, described as regions A-D, include *rpoB*, *rpoC*, other ribosomal genes, rPE and PPE family members.

For all other regions, we assigned a consensus base call to each position. A confident base call of a single base (A,C,G,T) was assigned, based on mapped high-quality bases (as above), provided

(i) it was not mixed (‘M’), as defined above;
(ii) sequence depth exceeded a prespecified cutoff of 10,
(iii) one base accounted for >90% of the pileup, a commonly used rule^2^.

All other bases were recorded as uncertain (‘N’).

#### Inter-lineage mixtures

We identified *M. tuberculosis* lineage using the published phylogeny^6^ and assessed inter-lineage mixtures of two, or more than two, samples using F2 and F47 statistics, as described^7^.

#### Computation of single nucleotide variation (SNV)

We compared sequences pairwise, considering a SNV present if a base (A,C,G,T) was confidently called in one sequence, and a different base confidently called in the other sequence. A SNV was counted at a given position if the two sequences differed in confident (A,C,G,T) consensus calls; it was not considered to be present if one or both consensus calls were uncertain (N) or mixed (M).

#### Detection of mixed sequences and the MixPORE algorithm

Suppose we have a NGS sequence *s_t_*, which we wish to test for mixtures of samples. *s_t_* forms part of a large collection s of *M. tuberculosis* sequences from a prospective sequencing program which densely samples TB isolations in a geographic region. We wish to determine whether *s_t_*_._ is likely to represent a mixture of other members of the collection *s* derived from different individuals. M sites are those where multiple bases maps to a single position (Fig. S1A,B). Mixed positions are ignored in SNV distance computations, so mixed samples appear highly similar or identical to their ancestors using SNV distance estimates (Fig S1C).

Biologically, *bona fide* mixed positions are likely to exist at positions of recent evolution (PORE) because (i) TB evolution is ongoing, and (ii) mixed samples commonly represent co-infection with, or co-evolution of, related samples^8^ (Fig S2A-C). Suppose we know *n* other sequences s_1,_ s_2_, .. s_n_ lie within some SNV threshold (say, 5, 20 , 50 or 100 SNV) of consensus base called data from *s_t_* (Fig 2B,C). If *s_t_* is in fact a mixture, the n sequences identified are expected to include close neighbours similar to both components of mixture (Fig. S1C). The *k* positions varying between the set of related sequences {*s_t_*, s_1,_ s_2_, .. s_n_} we denote as set P; in settings with dense sampling of TB isolates from the community, we expect P to includes PORE (Figure S2A).

The MixPORE algorithm tests the hypothesis that in the sequence *s_t ,_* M-sites occur at significantly higher frequency in the positions of recent evolution P relative to all other positions (P’). We tested this hypothesis by

(i) estimating the expected frequency *f*_P’_ of mixed bases in the positions P from as the number of M sites in P’/number of positions in P’. The size of the set P’ is large (~ 4 million bases) relative to the size of P, which was typically less than 100 sites.
(ii) comparing the observed number of M sites in P with the estimated expection *f*_P’_ using a binomial test. We used the python 3.7 scipy.stats.binom_test function to do this process, which is illustrated in Figure S2D.

#### Simulation of TB transmission in England

We simulated phylogenies representing local outbreaks in which 50 sequences originated from a single ancestor. Simulation used a random birth-death process ^9^, with birth:death ratio of 1:0.7, implemented in python (Fig. S2A). We simulated true nucleotide sequences of 10,000 nt, compatible with these phylogenies, using *pyvolve* ^10^ (Fig S2B) with evolution in up to 0.5% of nucleotides. We simulated observed consensus sequences in which (i) a random selection of 1% of bases are called uncertain (N); (ii) a random selection of either 0.01% or 0.1% bases are called mixed (M), rates corresponding to the error models observed for sequences obtained using HiSeq and MiSeq/NextSeq technology (see Supplementary Results). We generated mixed sequences from two sequences, randomly selected from the fifty, generating mixed bases (M) called at positions of variation (Fig S2C). For each phylogeny, we applied the MixPORE algorithm to each sequence, comparing observed with expected results.

#### Relationship between total mixed base numbers and those found by MixPORE

We studied the relationship between the number of mixed bases detected by MixPORE (M_mixPORE_), and the total number of mixed bases identified by cross-genome per-base binomial testing (M_total_). To do so, we stratified the data by sequencing technology (HiSeq *vs.* MiSeq) and modelled

M_total_ = k M_mixPORE_ +c, where c is M_total_ when M_mixPORE_ = 0 and k is the constant by which M_total_ increases for every increase in M_mixPORE_. Because of influential outliers, we used quantile regression to fit models using the R quantreg package.

#### Risk factors for mixed infection

To examine risk factors for mixed infection, we applied the MixPORE algorithm using a range of SNV thresholds to identify related samples and thence identify PORE. Analyses were run on the whole data set separately for 5, 20, 50 and 100 SNV thresholds. First, we classified samples as mixed (defined as at least one MRE-site) or not mixed. Next, we assessed the relationship between mixture status and a series of predictors, ascertained by questionnaire during routine surveillance of TB in England and stored in the PHE Enhanced TB Surveillance System, using univariate and multivariate logistic regression with the R 3.31 glm command.

#### Mixed bases and isolation of subsequent similar samples

To determine whether the detection of mixed bases might be associated with TB transmission, we applied MixPORE to the samples in the order the samples were received, only identifying neighbours among samples (i) received before the sample of interest and (ii) from a different individual. Then, we computed the time to isolation of a similar sample with properties compatible with transmission. We considered samples compatible with transmission if they were (i) 0-3 SNV of the first case (ii) were isolated after the first case, but within a year of it, and (iii) were isolated from a different person. A range of sensitivity analyses were done. In one, samples with more than one mixed base, as detected by MixPORE, were excluded, and (since mixed samples may underestimate divergence), for mixed samples, only subsequent similar samples within a penalised cutoff of 2 SNV were considered possible transmissions. Cox Proportional Hazards and Kaplan-Meier time to similar isolation analyses, censored at 1 year, and stratified by mixture status, were conducted using the R survfit package.

#### Software implementation

The Public Health England bioinformatics pipeline used for TB processing is freely available at <https://github.com/oxfordmmm/CompassCompact>. Code used to parse VCF files and perform per-base binomial testing is at <https://github.com/davidhwyllie/VCFMIX>. Fasta files containing consensus base calls, or IUPAC codes consisting of high-confidence mixtures, were generated using the VCFMIX fastaMixtureMarker module. These fasta files were loaded into a server implementing the mixture detection computation; software is available at <https://github.com/davidhwyllie/findNeighbour3>. The software comprises a pure python server based bacterial relatedness monitoring solution, accessible via RESTful endpoints. The mixture assessment process is provided by the */api/v2/multiple_alignment* endpoint. A python client is also provided. Performance characteristics of the software and technical details will be presented separately. Code performing simulations is in the *make_* and *run_simulation_clinical_paper.py* scripts.

### Supplementary Results

#### Estimates of mixed base numbers differ markedly depending on sequencing technology

The classification of mixed *vs.* unmixed samples requires an understanding of how many apparently mixed bases result from technical factors (false positive mixed base assignations), as opposed to reflecting biological mixtures (true positive mixed base assignations).

Because sequencing technology is an important determinant of sequencing quality, we compared the results of sequencing and reference mapping (see Methods) of ten reference Mycobacterial DNA preparations (provided by National Institute for Public Health and the Environment (RIVM), Netherlands, as part of a quality control distribution) using three different Illumina sequencing instruments in use in our institution (Table S1). Twenty-nine sequences were available; one sequence failed, and was not repeated (Table S1). The high-quality read mapped depths achieved differed between instruments used (Figure S3). A population of mixed positions (M-sites) was evident in all samples (Figure 2), a proportion of which occurred in clusters along the chromosome (Figure 2, yellow bars). M-sites, N-sites, and sequence depths with different sequencing technologies are shown in Fig. S4A-C, respectively.

#### Statistical testing for mixed sequences

Because both minor variant frequency, and the number of M-sites, differ markedly by sequencing technology, we developed a new approach (MixPORE) to mixture detection, which is intended to be insensitive to these confounders.

This approach relies on comparing the number of M-positions located within positions of recent evolution (PORE) with the number of Msites located in other parts of the genome (see Methods, and Fig S2D). We assessed the performance of MixPORE in the detection of mixed sequences in a simulation, designed to represent the scenario currently existing in England. In this simulation, *M. tuberculosis* transmission is ongoing, with diversification from common ancestor(s), while technical factors cause the reporting of uncertain calls (Ns) and false positive mixed calls (Ms) at rates similar to those observed with HiSeq and MiSeq sequencing in our laboratories.

MixPORE assesses mixtures in positions of recent evolutionary change, and so it cannot determine PORE unless at least two similar samples are present. Additionally, in this work we only considered samples from other patients. Analysis of 100,000 sequences in 2,000 randomly generated phylogenies shows MixPORE has near perfect sensitivity and specificity when determining mixtures in samples with at least two close neighbours (Fig. S5). However, if most samples do not have close neighbours sampled – as is the case in over 80% of samples in the simulations run - the overall sensitivity of mixture detection is low (Fig. S5). Simulations also indicate that the sensitivity of mixture detection is not strongly influenced by MiSeq *vs.* HiSeq sequencing technology (Fig. S6).

### Supplementary Data

#### Supplementary Data D1

This describes the results of various sensitivity analyses performed, and is provided as a separate, self documented Excel workbook containing model coefficients from range of analyses. To examine risk factors for mixed infection, we applied the MixPORE algorithm using a range of SNV cutoffs c to identify similar samples and thence identify PORE. Analyses were run separately for c= 5, 20, 50 and 100. First we classified samples as mixed (defined as at least one mixed based identified using MixPORE) or not mixed. Next, we assessed the relationship between mixture status and a series of predictors, ascertained during routine surveillance of TB in England and stored in the PHE Enhanced TB Surveillance System, and the likelihood of a sample being mixed using univariate and multivariate logistic regression using the R glm command using R 3.3.1.

### Supplementary Table

#### Table S1 Samples investigated

| **External ID** | **Centre 2 Specimen Id** | **Centre 1 Specimen Id** | **MiSeq Derived Sequence Identifier** | **NextSeq Derived Sequence identifier** | **HiSeq derived sequence identifier** |
| --- | --- | --- | --- | --- | --- |
| NGS1 | H184440695 | 18.624138 | f32a6c5a-503f-4c50-be10-5b98ed26f710 | df109200-b074-4480-99c2-670510fa7120 | 027cdb24-e8d5-421a-a808-6cbf1667baa1 |
| NGS2 | H184440696 | 18.624139 | 79a91c9d-71f7-40bd-a900-e310aeef2cfe | f91f994d-02dc-45f9-b463-6ad2f3308b69 | 27d2f2cf-1d20-4e85-87c1-c8d440ec8b85 |
| NGS3 | H184440697 | 18.624140 | f70514c8-c3d7-4270-a0ce-b7642d7c6aa7 | 23365501-7e6c-44bd-b30f-b1ac2c78a6c0 | 48399e35-4e1f-49b4-b7cf-7e25281d3f8a |
| NGS4 | H184440698 | 18.624141 | c7591a7d-3897-4446-abcc-ae5b19656b14 | Not determined | e504cf35-f231-42e5-a08a-899e1e27dc64 |
| NGS5 | H184440699 | 18.624142 | 6cfa34f5-0c74-4cf9-bc2e-4030b93f93f8 | a31647d9-25d6-4526-b46e-b09af4b42bff | f27f3402-c581-4803-8cb4-fe2aa2089b80 |
| NGS6 | H184440700 | 18.624143 | c9de7445-e6f5-49e3-ac24-bfcbec4a4db0 | 8a2d3d1c-da78-4462-9fd6-81b201954bea | 03532a28-a095-4d0b-89e8-cfe1b47639b0 |
| NGS7 | H184440701 | 18.624144 | a37054d6-95e4-4428-b578-919d73b7e562 | fa88a87f-3f0d-4c8b-94d5-25e5e79f8dd0 | ae3a6dde-1c06-4016-a0e3-0a7d1be6c36f |
| NGS8 | H184440702 | 18.624145 | 36e35aa7-ac77-4442-8d3d-0ef512d2f9bd | ccf10511-9f08-4138-9380-5e71361f2014 | 26f75f4f-d921-460a-87d3-675fa623c499 |
| NGS9 | H184440703 | 18.624146 | 4bb1e260-ce12-43eb-9679-548f08f983fd | 5d8a0c37-6b68-4438-91f5-cea93743b8d8 | 5cf243a1-182e-4fe7-acd9-1f1aae61aef1 |
| NGS10 | H184440704 | 18.624147 | 193e4ef0-03a6-4232-8d62-fa8e4187229a | d7f63bf9-b386-4748-9dde-329983c3fe9a | e8888e07-00c7-41ae-a701-2b3884bfe738 |

#### Table S1 Legend

Sequencing identifiers of the ten samples used for comparison of sequencing technologies.

### Supplementary Figures

#### Figure S1 Mixed bases cause systematic underestimation of pairwise SNV

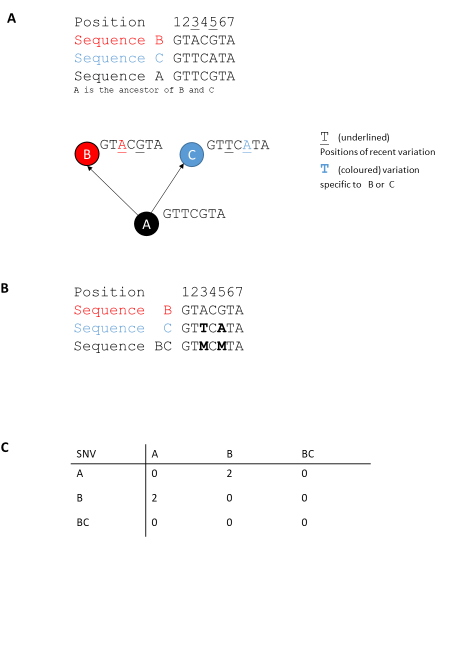

#### Figure S2 Illustration of MIX-PORE algorithm for mixture detection

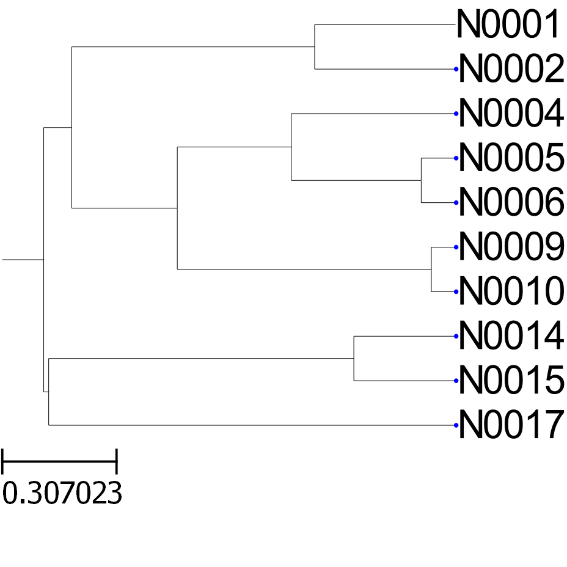

**A Phylogeny**

N0001 GGGGGGGGGTCGCCGAATAGGTT

N0002 GGGGGGGGGTGCCCAATTTGGCT

N0004 GGGGGGGGGTGTGCGAGAACGCG

N0005 GGGAGGGGGTGGCCGAGTAGCCC

N0006 GGGGGGGGGTGGCCGAGTAGCAC

N0009 GGGGGGGGGTCTCCCCTAGCGCC

N0010 GGGGGGGGGTCTCCCCTACCGCC

N0014 GGGGGGGGGGCATTGTCGACGGT

N0015 GGGGGGGGGCAACTGACGACGGT

N0017 GGGGGGGGGAGATGACTCTCCCT

**B True sequences**

```
N0001-4* GGGMGGGGGTMMMCGAMMAMGMM
N0002    GGGGGGGGGTGCCCAATTTGGCT
N0004    GGGAGGGGGTGTGCGAGAACGCG
N0005    GGGGGGGGGTGGCCGAGTAGCCC
N0006    GGGGGGGGGTGGCCGAGTAGCAC
N0009    GGGGGGGGGTCTCCCCTAGCGCN
N0010    GGGGGGGGGTCTCCCCTACCGCC
N0014    GGGGGGGGGGCATTGTCGACGGT
N0015    GGGGGGGGGNAACTGACGACGGT
N0017    GGGGGGGGGAGATGACTCTCCCT
*N0001-4 the sequence called from
N0001 mixed with N0004
```

**C Observed consensus**

**sequences**

**D Algorithm determining whether a sample is mixed**

1. Identify sequences similar to the sample of interest, in this case N0001-4, using SNV estimates

- For N0001-4, all samples in C are within 10 SNV, and each is from a different patient.

2. Define the positions of variation between these samples, reflecting Positions of Recent Evolution (Pore)

- The positions varying between the sequences are boxed in Panel C

```
3. Determine the number of M calls in the Positions of recent evolution, vs. the rest of the sequence
                Rest of the sequence             Positions of recent evolution
                GGGMGGGGG                        TMMMCGAMMAMGMM
Number of Ms    1                                8
4. Determine estimate of expected probability of observing M calls based on rest of the sequences
Expected p = 1/ 9
5. Determine observed number of M calls in the positions of recent evolution
Observed 8 in 14 positions
6. Compare observed frequency with expected frequency using binomial test.
```

ancestral

Recently

evolved

#### Figure S3 Characteristics of sequencing by MiSeq vs. HiSeq technology

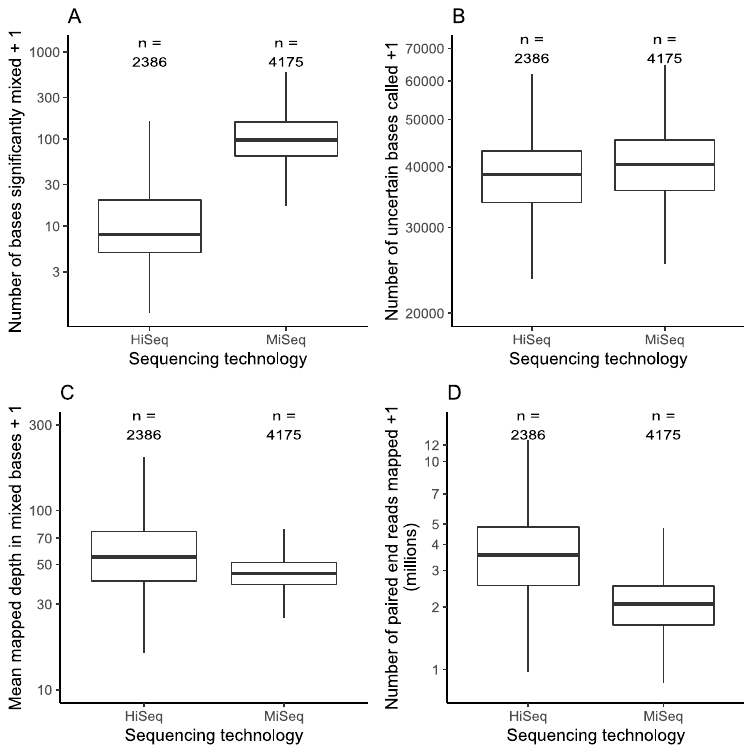

#### Figure S4 Sequenced depths with different sequencing technologies

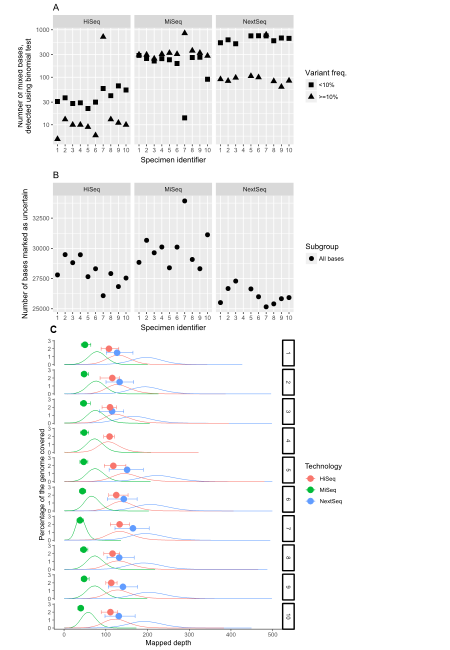

#### Figure S5 Detection of mixed samples by MixPORE: assessment by simulation

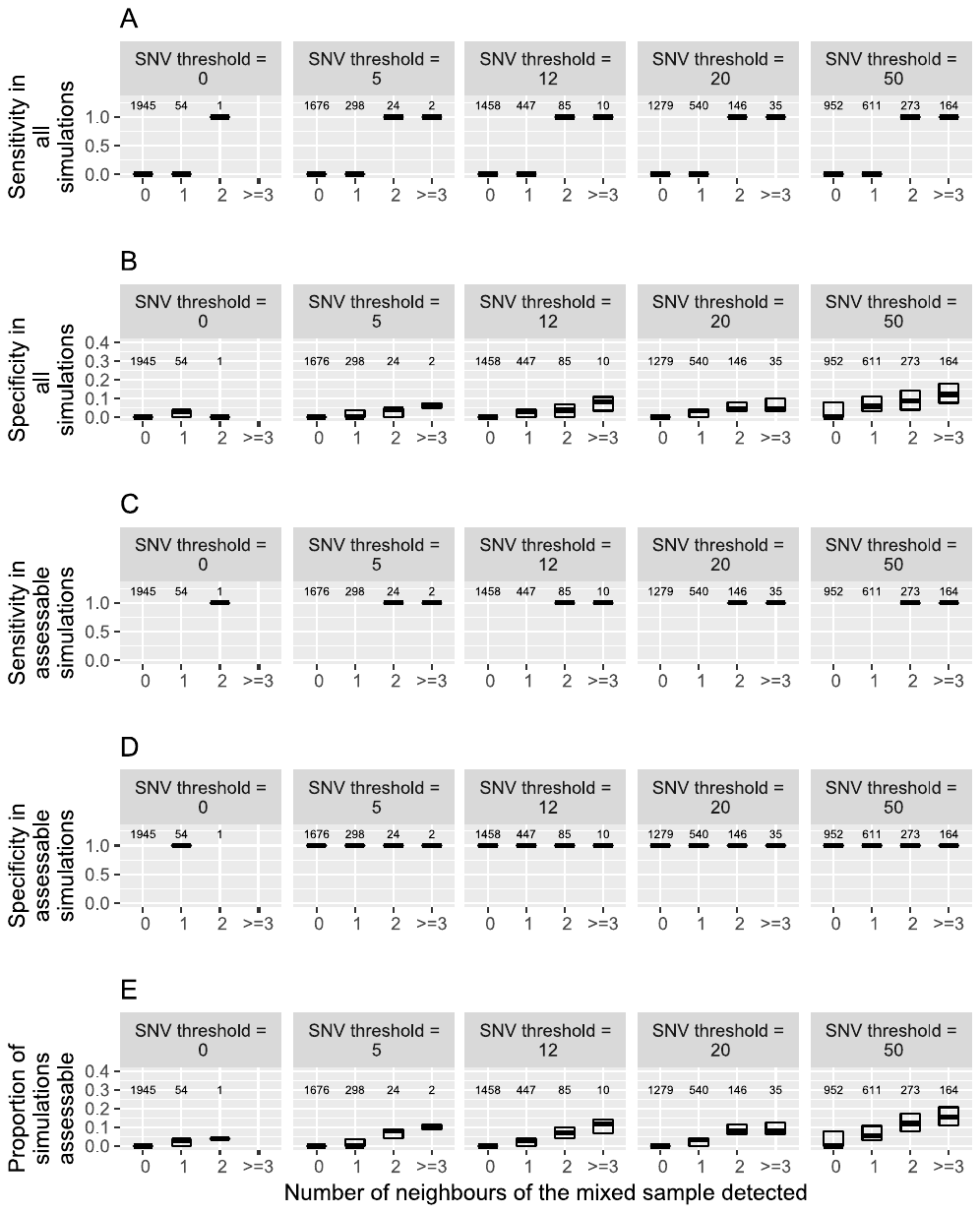

#### Figure S6 Impact of sequencing platform on mixture detection: assessment by simulation

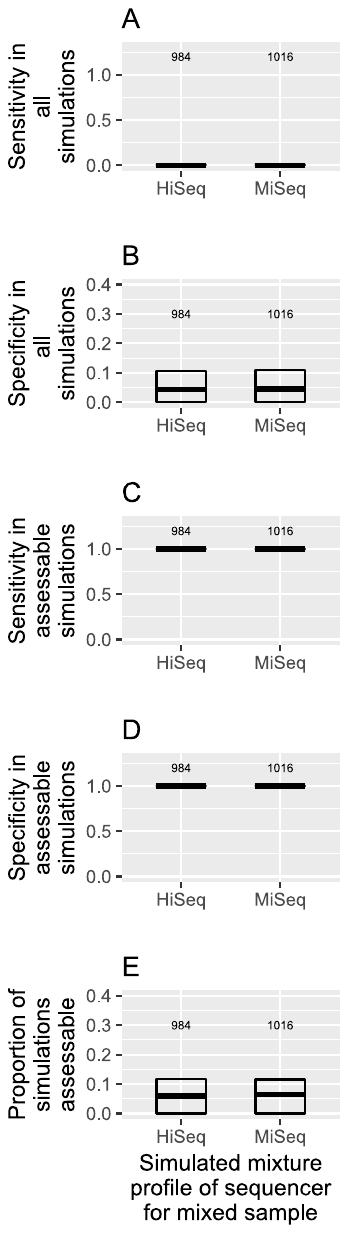

#### Figure S7 Relationship between total mixed base detection and that by MixPORE: 5 SNV threshold used by MixPORE

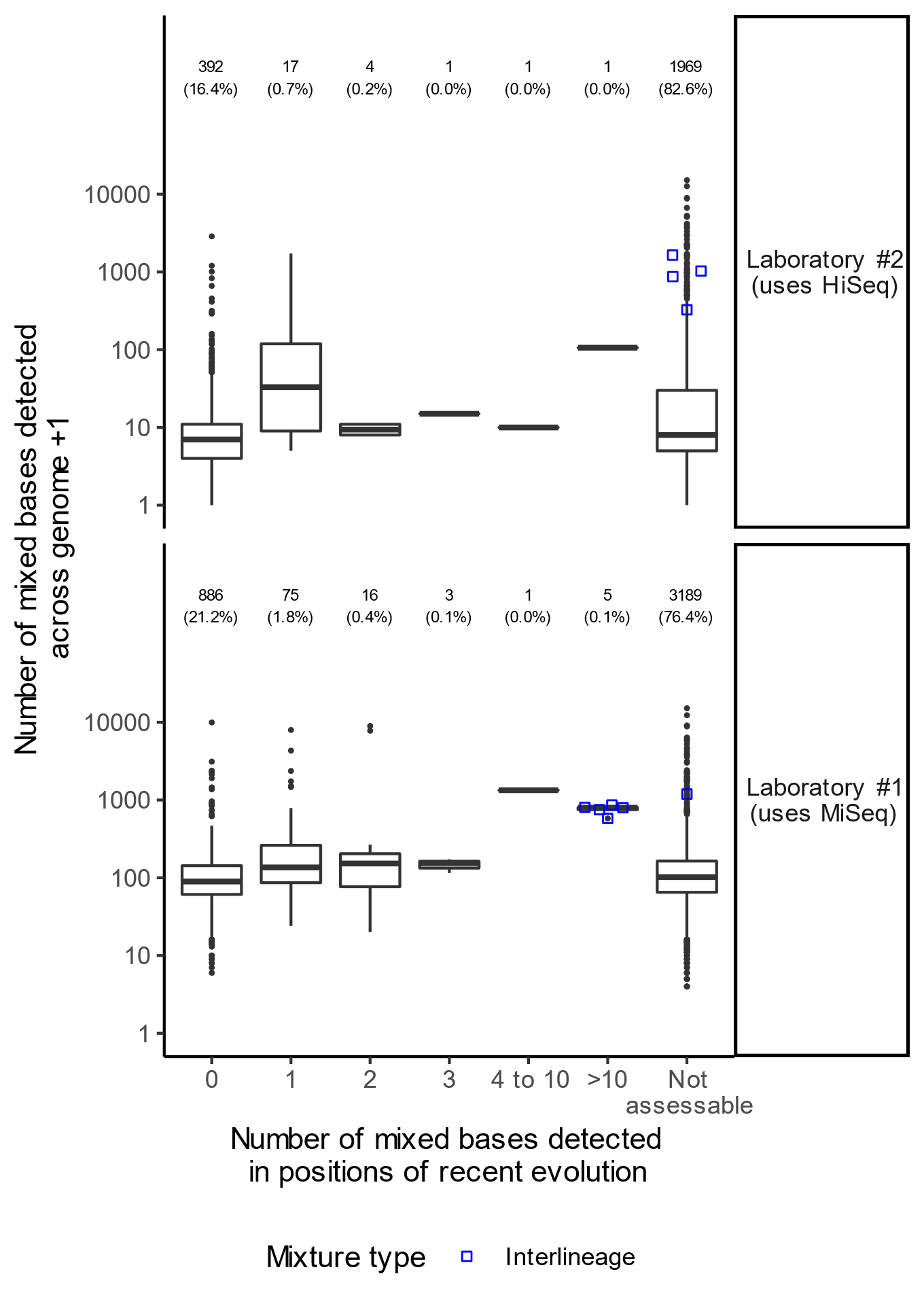

#### Figure S8 Relationship between total mixed base detection and that by MixPORE: 20 SNV threshold used by MixPORE

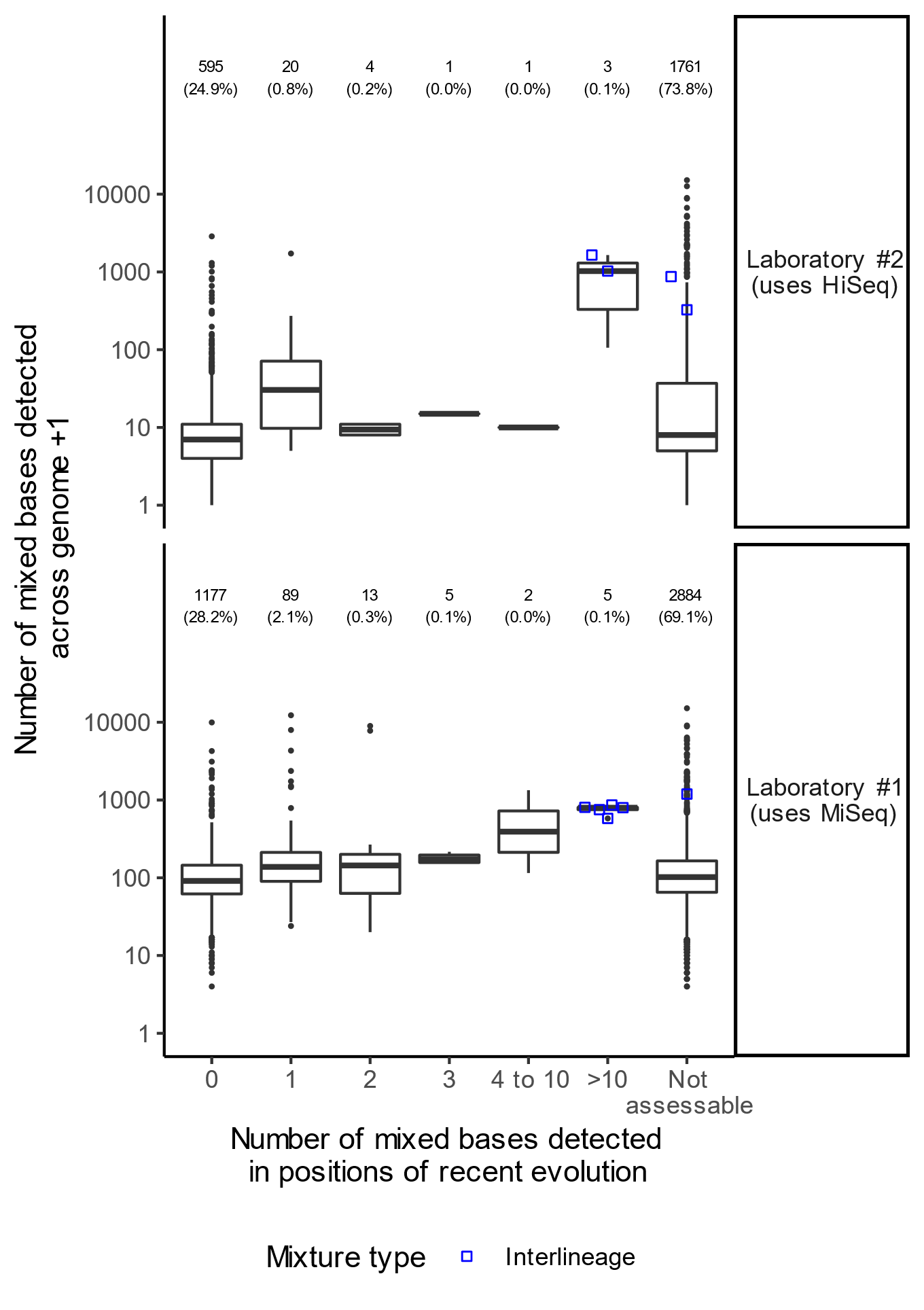

#### Figure S9 Relationship between total mixed base detection and that by MixPORE: 50 SNV threshold used by MixPORE

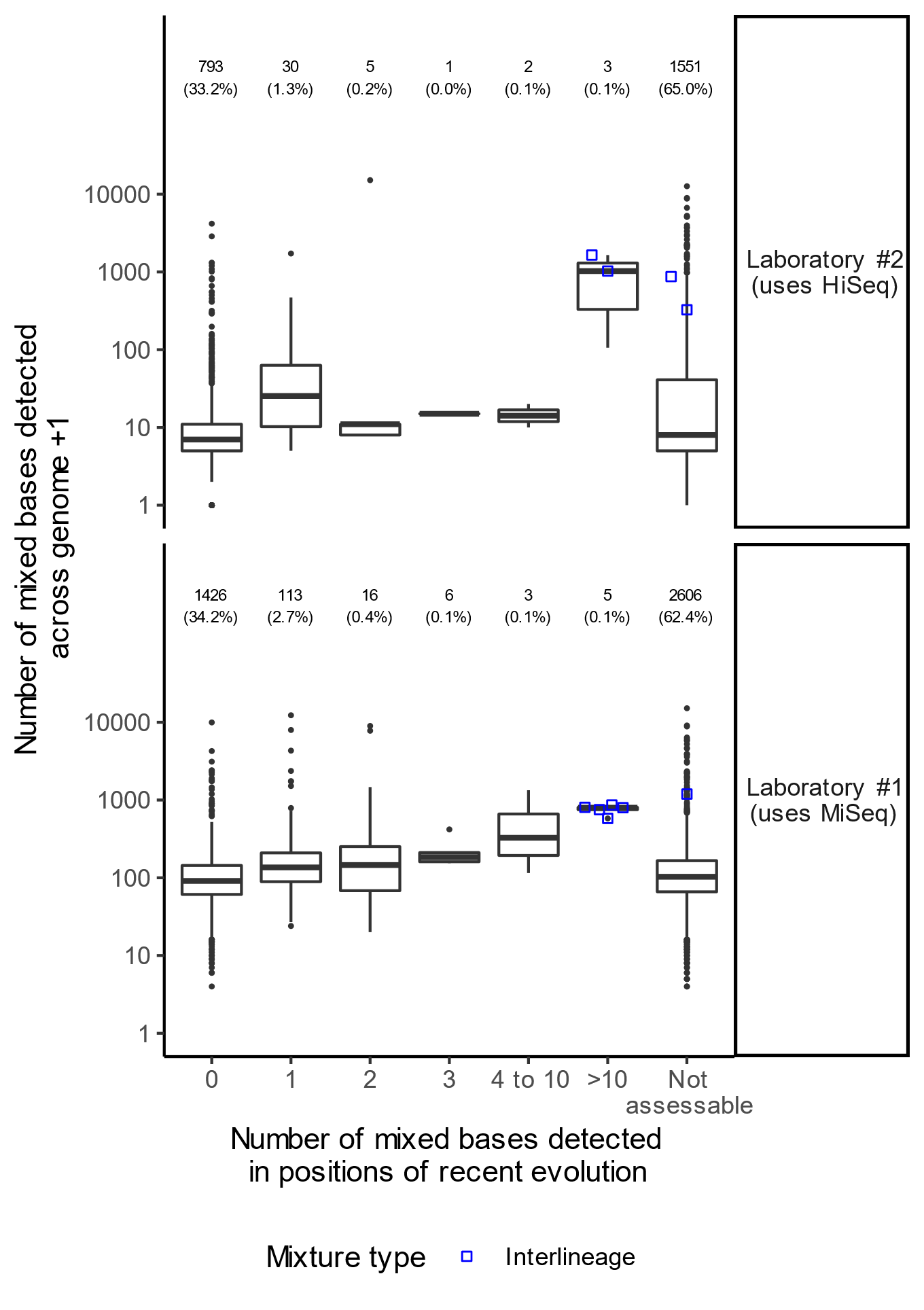

#### Figure Legends

##### Figure S1 Mixed bases cause systematic underestimation of pairwise SNV

A: Three sequences, each of seven nucleotides, are illustrated. A is the ancestor of B and C. Positions of recent evolution are underlined; such positions include the variation between B and C.

B: Sequence BC is the consensus sequence called from a mixture of B and C. M denotes a mixed base.

C: A distance matrix formed between the consensus sequences in of sequences A,B and BC.

##### Figure S2 Illustration of MIX-PORE algorithm for mixture detection

A: Consider a phylogeny of *M. tuberculosis* samples, originating from a common ancestor, such as that shown in the simulated phylogeny in A. These sequences comprise the sequence of the ancestor, here illustrated as being all guanine nucleotides (B), and recently evolved change. If a mixture is present, after mapping, multiple bases will map to a single position at positions of recent evolution (M sites) (C). M sites may also occur for technical reasons, but these will be scattered across the genome, not overrepresented in positions of recent evolution. This forms the basis for an algorithm (D) comparing the representation of Ms in these two populations of bases.

##### Figure S3 Characteristics of sequencing by MiSeq vs. HiSeq technology

Characteristics of sequences obtained from MiSeq vs. HiSeq technology, depicted as box and whisker plots. The numbers refer to the number of sequences sequenced by each technology. The number of bases detected as mixed (‘M’) differed with a median of 7 vs. 99 bases, respectively. The high quality read depth also differed between technologies (median 44 vs. 54), as did the number of read pairs (2.0 vs. 3.5 x106) and numbers of uncertain bases (‘N’) called (41,092 vs. 39,005) (p < 10-8 for all four comparisons).

##### Figure S4 Sequenced depths with different sequencing technologies

Ten reference *M. tuberculosis* isolates, provided as part of an External Quality Assessment panel, were sequenced using three different Illumina technologies. Reads were mapped to the H37Rv v2 reference genome.

A: number of mixed bases, detected by binomial testing, for each of ten samples processed using three different sequencing technologies. Y axis is a logarithmic scale.

B: number of bases marked as uncertain by the sequencing pipeline, for each of ten samples processed using three different sequencing technologies.

C: The sequencing depth obtained at all bases (kernel density estimates, smooth lines) and a ‘M sites’ identified as mixed using a Binomial test (median shown as a dot; 5^th^ and 95^th^ centiles as whiskers) are shown for each sequence.

##### Figure S5 Detection of mixed samples by MixPORE: assessment by simulation

2,000 simulations were performed in which artificial phylogenies were generated, a single mixed sequence produced from simulated sequences, and errors introduced corresponding to known error profiles of observed sequencing pathways (see Methods). Half the samples had error profiles for MiSeq and half for HiSeq machines introduced. For each phylogeny, we determined (a) the number of neighbours (determined by SNV threshold: SNV thresholds of 0,5,12,20, and 50 were examined; in this simulation, more than 50 SNV were rarely present between simulated sequences) of the mixed sequence (b) whether the mixed sequence was recorded as mixed (c) whether other sequences were recorded as mixed. Sensitivity and specificity were computed for (a) assessable simulations, which were those in which the mixed sample had at least two neighbours and (b) all simulations. Numbers indicate the total number of simulations in each category.

##### Figure S6 Impact of sequencing platform on mixture detection: assessment by simulation

2,000 simulations were performed in which artificial phylogenies were generated, a single mixed sequence produced from simulated sequences, and errors introduced corresponding to known error profiles of observed sequencing pathways (see Methods). Half the samples had error profiles for MiSeq and half for HiSeq machines introduced. For each phylogeny, we determined (a) the number of neighbours (determined by a SNV threshold of 50) of the mixed sequence (b) whether the mixed sequence was recorded as mixed (c) whether other sequences were recorded as mixed. Sensitivity and specificity were computed for (a) assessable simulations, which were those in which the mixed sample had at least two neighbours and (b) all simulations. Numbers indicate the total number of simulations in each category. Data are presented stratified by the sequencing technology used for the mixed samples. There are no significant differences for any of the parameters plotted (Mann-Whitney tests, p > 0.1).

##### Figure S7,8,9 Relationship between total mixed base detection and that by MixPORE

Relationship between the numbers of mixed bases (M-sites) detected using per-base examination (y-axis) and that detected by MixPORE at positions of recent evolution (MRE-sites) (x-axis). The analysis is stratified according to whether the sample was analysed by MiSeq or HiSeq based processes. Numbers show number (percent of total for MiSeq/HiSeq) of samples in each group. Large circles indicate inter-lineage mixtures, detected by F2 statistic computation as described^7^. These figure shows results when a range of cutoffs are used to select neighbours for the MixPORE algorithm.
